# Biodiversity dynamics with complex genotype-to-phenotype architecture in multilayer networks

**DOI:** 10.64898/2026.03.23.713274

**Authors:** Carlos J. Melián, Cecilia S. Andreazzi, Julia Astegiano, Víctor M. Eguíluz, Francisco Encinas-Viso, Philine G. D. Feulner, Luis J. Gilarranz, Paulo R. Guimarães, Ruben Heleno, Weini Huang, François Massol, Jordi Moya-Laraño, Jelena Pantel, Cas Retel, Pooja Singh, Ali R. Vahdati, Blake Matthews

## Abstract

2

The genotype-to-phenotype architecture (GPA), defined by complex interactions such as pleiotropy, epistasis, and regulatory control, is a fundamental yet often overlooked driver of biodiversity dynamics. While empirical evidence suggests that traits mediating species interactions (biotic) and environmental responses (abiotic) are frequently correlated, most eco-evolutionary theories treat these traits as independent, leaving a gap in our understanding of how genomic architecture influences community-level outcomes. In this study, we contrast two distinct GPAs, modular (independent trait evolution) and correlated (integrated trait evolution), within a spatially explicit multilayer network framework. We evaluate their impact on biodiversity across varying regimes of selection, migration, and biotic and environmental filtering. Our results reveal a hierarchy of drivers: selection strength dictates the absolute magnitude of the architectural effect, while migration and context-dependent biotic and abiotic effects determine which architecture yields a diversity advantage. Correlated GPAs enhance species coexistence and diversity in low-migration landscapes characterized by strong selection and moderate, balanced biotic and abiotic pressures. In these contexts, trait integration serves as a buffer against selective noise. Conversely, modular GPAs support higher diversity under high migration and strong biotic interactions, where the decoupling of trait modules provides the adaptive flexibility necessary to navigate spatially conflicting selective pressures. Our findings demonstrate that genomic architecture acts as a critical filter for environmental perturbations. Integrating complex GPAs into multispecies models is essential for quantifying the co-evolutionary feedbacks among traits, population adaptation, and species persistence. Our framework provides a path for predicting how biodiversity emerges and persists across biological scales, from genomics to communities and food webs, under the accelerating pressures of global change.

**Conclusions:**

1. We integrate trait architecture to spatial biodiversity to show biodiversity patterns are not merely products of ecological interactions, but are fundamentally constrained by Genotype-to-Phenotype Architecture (GPA). By linking GPA to biodiversity we show the interplay between the complexity of an organism and community structure in determining diversity patterns.
2. The hierarchy of Eco-Evolutionary Drivers: We establish a new conceptual hierarchy where selection strength acts as the fundamental governor of architectural impact, while the specific architecture predicting higher diversity (Correlational vs. Modular) is dictated by the interplay of migration scales and context-dependent biotic and abiotic dynamics.
3. Selection-Migration contingency for coexistence: We provide a new hypothesis for species coexistence: Correlational selection serves as a stabilizing force under dispersal limitation, whereas Modular trait architecture provides the adaptive flexibility to maintain diversity in high-migration, spatially heterogeneous landscapes.
4. Adaptive decoupling as a diversity engine: We propose that trait modularity functions as a “buffer” against extinction by decoupling phenotypic responses. This allows populations to navigate conflicting selective pressures, effectively facilitating evolutionary rescue in complex biotic environments.
5. Methodological framework for empirical inference: To bridge the gap between theory and data, we provide a novel likelihood-based framework. This enables researchers to infer latent trait architectures from population genomic samplings, turning GPA from a theoretical construct into a measurable sampling variable in natural populations.
6. We define a new roadmap for the next generation of eco-evolutionary modeling. By identifying the gaps between existing simulation engines, we provide a conceptual “blueprint” for a digital ecosystem that fully integrates complex genetic architecture with global bio-diversity dynamics.

## 3 Introduction

Biodiversity is declining globally at unprecedented rates, driven by intensive land use and growth-oriented economies (Brondizio et al., 2019). This erosion of species richness does more than simplify ecosystems; it degrades the genetic and trait variation that underpins ecosystem resilience and functional complementarity (Loreau et al., 2001; Weisser et al., 2017; Fanin et al., 2018). Biodiversity loss also deteriorates ecosystem resilience by reducing the number of species and phenotypes that can resist future environmental changes (Calow, 1987; McGill et al., 2006; Violle et al., 2007; D íaz et al., 2016). While we recognize that species’ responses to such change are shaped by biotic and abiotic traits, our predictive capacity is currently stalled by a “complexity gap”: the disparity between the nonlinear dynamics of genotype-to-phenotype maps (GPM) and the simplified models of trait dynamics typically used in ecological forecasting (de Villemereuil, 2018; Milocco and Salazar-Ciudad, 2022). To bridge this gap, we must move beyond treating traits in isolation. Critical adaptive responses, such as drought resistance or foraging efficiency, are governed by a dense architecture of epistasis and pleiotropy. Understanding this architecture is essential for forecasting whether species will adapt to climate change, invade new habitats, or succumb to extinction. Despite a consolidation of trait-based approaches in ecology and evolution (Cavender-Bares and Wilczek, 2003; Carmona et al., 2016; Rolland et al., 2023), the link between complex trait architecture and the dynamics of multi-species communities remains largely undeveloped (Bailey et al., 2009).

Ecological approaches have embraced the connection between biotic and abiotic traits by considering how traits are structured within and among populations (Laughlin and Messier, 2015). Trait-based approaches have revealed key mechanisms of biodiversity dynamics (Tilman et al., 1997; Carmona et al., 2016; Violle et al., 2007; McGill et al., 2006; Enquist et al., 2015), including species turnover (Bannar-Martin et al., 2018), resistance and recovery (Cunillera-Montcus í et al., 2021), ecological niches and species distributions (Soberón and Peterson, 2005) and structural changes in species interaction (Maureaud et al., 2020; de Andreazzi et al., 2020). Evolutionary trait-based perspectives have also considered different traits and they are informative about how entire assemblages of interacting species respond to environmental change (Fussmann et al., 2007a). Many studies on trait evolution have shown how interaction traits modulate interaction networks by determining the interaction probabilities and their impact on fitness (Stang et al., 2007; Dehling et al., 2014; Schleuning et al., 2016). One of the main results of these approaches has shown that rapid changes in traits that govern interactions affect both interaction strength and network structure, including connectance and modularity (Loeuille and Loreau, 2005). Importantly, changes in the structure of direct and indirect interaction traits among species can alter selection pressures within communities, generating feedback between trait evolution and biodiversity dynamics in communities (Encinas-Viso et al., 2012; Nuismer et al., 2013; O. and C., 2016; de Andreazzi et al., 2020; Guimarães et al., 2017; Andreazzi et al., 2018).

There have also been many studies exploring the role of genetic architecture and genetic correlations, i.e., number of loci, linkage disequilibrium and ploidy on eco-evolutionary and coevolutionary dynamics (Endler, 1995; Nuismer and Doebeli, 2004; Tucker et al., 2012; Moya-Laraño et al., 2014). For example, the role of genetic architecture driving evolutionary rescue where rapid evolution prevents extinction (Yamamichi, 2022). It has been recently shown that highly modular trait architecture underlies the emergence of hundreds of species simultaneously as a predictor of an adaptive radiation, with the main hypothesis focusing on how recombining “only” within trait modules for a finite set of ecological traits drive speciation dynamics (Singh et al., 2025). Therefore, empirical patterns of weak genetic and ecological correlation across traits imply no selection constraints in trait associations to predict modular trait architecture and its consequences for speciation and adaptive radiations (Orkney et al., 2025). Overall, ecological, evolutionary and speciation studies point out to an important role of genetic architecture in rescuing populations, coevolutionary dynamics of interacting species and adaptive radiations. Yet, the contribution of complex GPM to biodiversity dynamics remains largely unexplored (Alberch, 1991; Salazar-Ciudad et al., 2021; Assis et al., 2020). Here, we refer to the Genotype-Phenotype Map as Genotype-to-Phenotype Architecture, GPA, by accounting for: 1) different classes of traits, i.e., biotic, abiotic and migration traits, and 2) different architectures, modular, all independent traits, vs. correlated accounting for interactions among traits. Genetic correlations between trait classes or the absence of correlations between biotic and abiotic traits might have potentially differential fitness effects and therefore many different routes for population dynamics, species interactions and biodiversity dynamics. The underlying Genotype-to-Phenotype Architecture (GPA), the structural “wiring” of how genetic correlations or their absence translate to phenotypic diversity, whether an architecture is Modular (independent trait classes) or Correlated (interacting trait classes) may fundamentally change how a population, communities and food webs traverse their fitness landscape (Orkney et al., 2025; Singh et al., 2025).

Interactions among genetic, trait and GPA components with biotic and abiotic traits can provide insights on how fast species respond to rapidly changing environments (Figures 1). Particularly, connecting GPAs to population dynamics can decipher how populations decline when confronted to multifarious perturbations, from habitat loss to climate change. The interactions between ecological and evolutionary processes involving GPAs might arise by multiple mechanisms operating at different levels (Govaert et al., 2019). Feedback can emerge between differences in phenotypes of interacting species (e.g. co-evolution), between population demography and trait evolution (e.g. evolution and demography with eco-evolutionary dynamics (Kokko and andrés, 2007)), and between organisms and abiotic conditions (e.g. evolution by niche construction (Matthews et al., 2014)). Currently, studies considering coevolutionary feedbacks have rarely accounted for different traits in response to biotic and abiotic selection pressures (Endler, 1995; Thompson et al., 2013; Guimarães et al., 2017; Weber et al., 2017; Terhorst et al., 2018). Existing trait approaches on biodiversity consider mostly biotic traits Kroumi and Lessard (2015); Assis et al. (2020) and simple genotype-to-phenotype maps, i.e., traits being influenced by multiple genes each having a small effect (but see (Singh et al., 2025; Milocco and Salazar-Ciudad, 2022)). Thus, exploring broader categories of architectures accounting for pleiotropy and epistasis can advance our understanding of how complex trait architecture respond to selection and migration (Melián et al., 2018; Yeaman, 2022).

**Figure 1.**
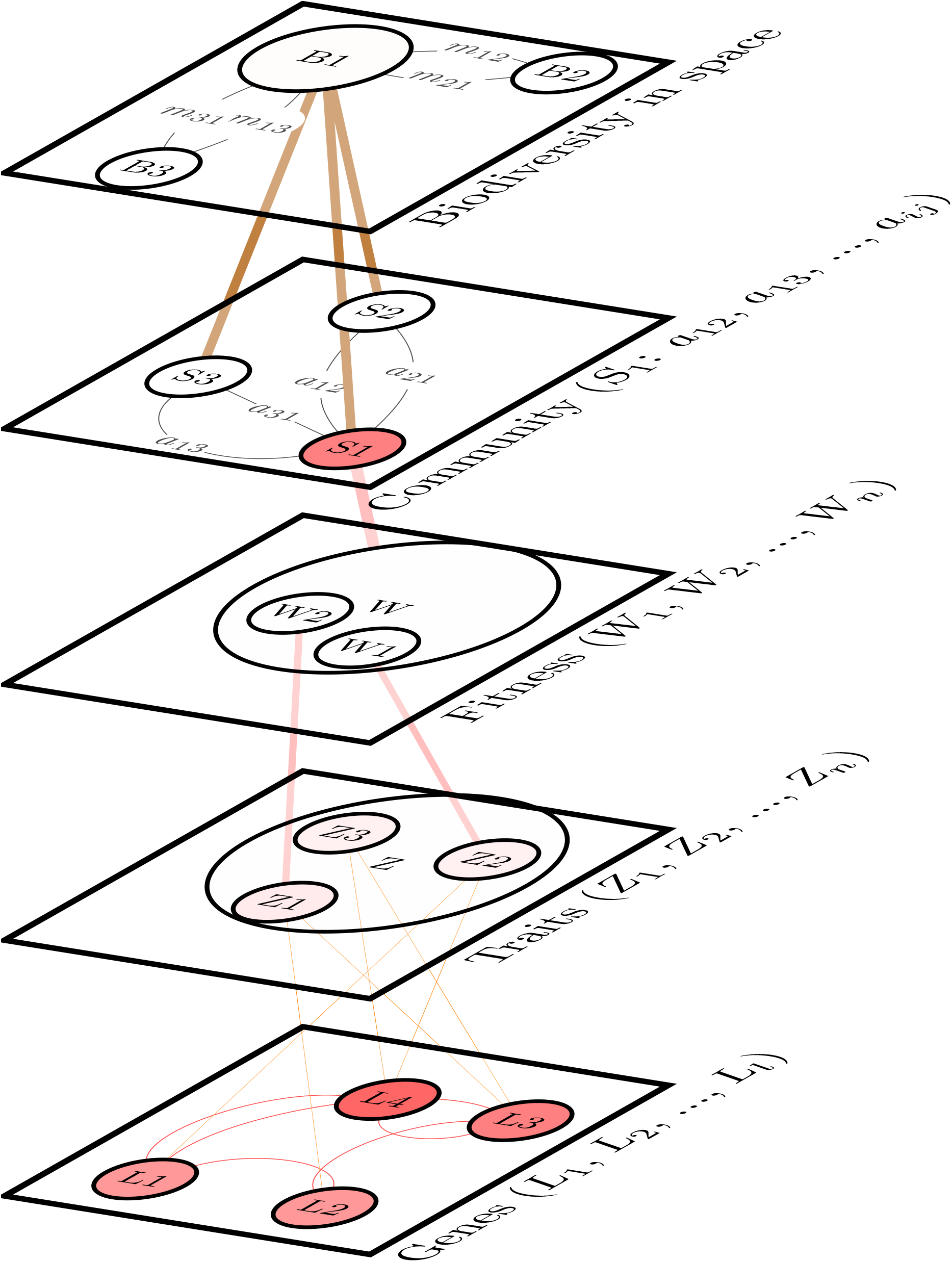
Multilayer network connecting complex genotype-to-phenotype architecture (GPA) to biodiversity dynamics: The modeling consists of five layers: Layer 1: Genes connect loci with epistasis and expression to layer 2, Traits. Genes are represented by loci, *l*1, *l*2, …, *lL*. Loci interactions represented with epistasis and expression (red links) and pleiotropy (orange links). Layer 2: traits for each individual represented as *z*1, *z*2, …, *zN*. Traits are compared with the biotic and abiotic states to compute the fitness (Layer 3). Fitness also accounts for the species interaction coefficients for each species, i.e., *S*1: *c*12, *c*13, …, *cij* (Layer 4). Population size for each site and total population size of a given species are then computed for many species (Layer 5). We compute diversity index to compare two GPAs predictions.

Many empirical studies suggest GPAs with traits jointly impacted by selection, but model systems containing large genetic and trait datasets to decipher the impact of GPAs on populations are only starting to emerge (Lynch, 2007; Svensson et al., 2021). Traits, like size, morphology, and those influencing responses to environmental and biotic conditions vary along continuous gradients. These traits can have heterogeneous effects on correlated traits and contribute differently to overall fitness. Traits are referred in some cases to as quantitative traits controlled by quantitative loci (QTLs) (Lynch and Walsh, 1998). Quantitative traits affecting fitness are often affected by multiple genes, i.e., polygenic background. If fitness traits are controlled by a large number of loci each with an infinitesimal small effect, then it becomes difficult to link selection to traits to understand adaptive changes at the molecular level (Barton, 2022). In this regard, many studies analyzing genomics data have shown few isolated loci having large phenotypic effect, a monogenic or oligogenic background (Enbody et al., 2022). Under this scenario, the genetic architecture contains complex traits composed of a small number of loci affecting fitness traits with varying strengths. Many combinatorial paths might exist in the GPA-to-fitness trait landscape, i.e., mono-, oligo-, poly-, and omni-genic, the last one referring to most or all genes expressed within a cell having an impact on the expression of a given trait, particularly, genes not directly impacting expression might be explaining more heritability than core genes, the ones with direct impact on expression, a process referred as “network pleiotropy” (Boyle et al., 2017).

There are many traits belonging to specific classes, i.e., biotic or abiotic traits, that can be correlated to each other, forming modules, and having a different contribution to fitness (Callebaut and Rasskin-Gutman, 2009; Wagner et al., 2007; Rubin et al., 2022). Trait correlations within a trait class, i.e., biotic traits, are often assumed to result from genetic correlations, but there might be a trait architecture as well. For example, trait correlations might also result from covariation among different sources of natural selection and interactions among the different trait’s classes, i.e., biotic and abiotic (Endler, 1995) but the interactions between biotic and abiotic traits might be based upon genetic correlations. If species are composed of many complex traits and traits respond to biotic and abiotic factors in many distinct ways, it can be important to understand the ecological and evolutionary processes underlying the GPAs to anticipate species response in rapidly changing ecosystems composed of many interacting species.

In this study, we introduce GPAs into a multilayer network framework to integrate complex traits within community and food-web dynamics. Multilayer networks offer many methods to facilitate the integration of ecological (Pilosof et al., 2017), developmental (Milocco and Salazar-Ciudad, 2022) and evolutionary processes (King and Wilson, 1975). We specifically contrast two GPAs modular and correlated, to identify the “master levers” predicting diversity patterns. Our results demonstrate that the architectural effect on diversity is not random; rather, it is governed by a strict hierarchy where selection strength determines the magnitude of the architectural impact, while migration and context-dependent state of biotic and abiotic traits dictate which architecture yields higher diversity. By treating GPAs as a complex filter for environmental noise, we provide a path to merge molecular systems biology with metacommunity and food web dynamics, offering a framework for predicting species’ fates in a rapidly changing world (Figures 1, 2). We connect GPAs to GWAS data and to trait distributions (Box 1) and diversity trajectories to foster the emergence of a digital ecosystem connecting complex trait architecture to biodiversity dynamics (Box 2).

**Figure 2.**
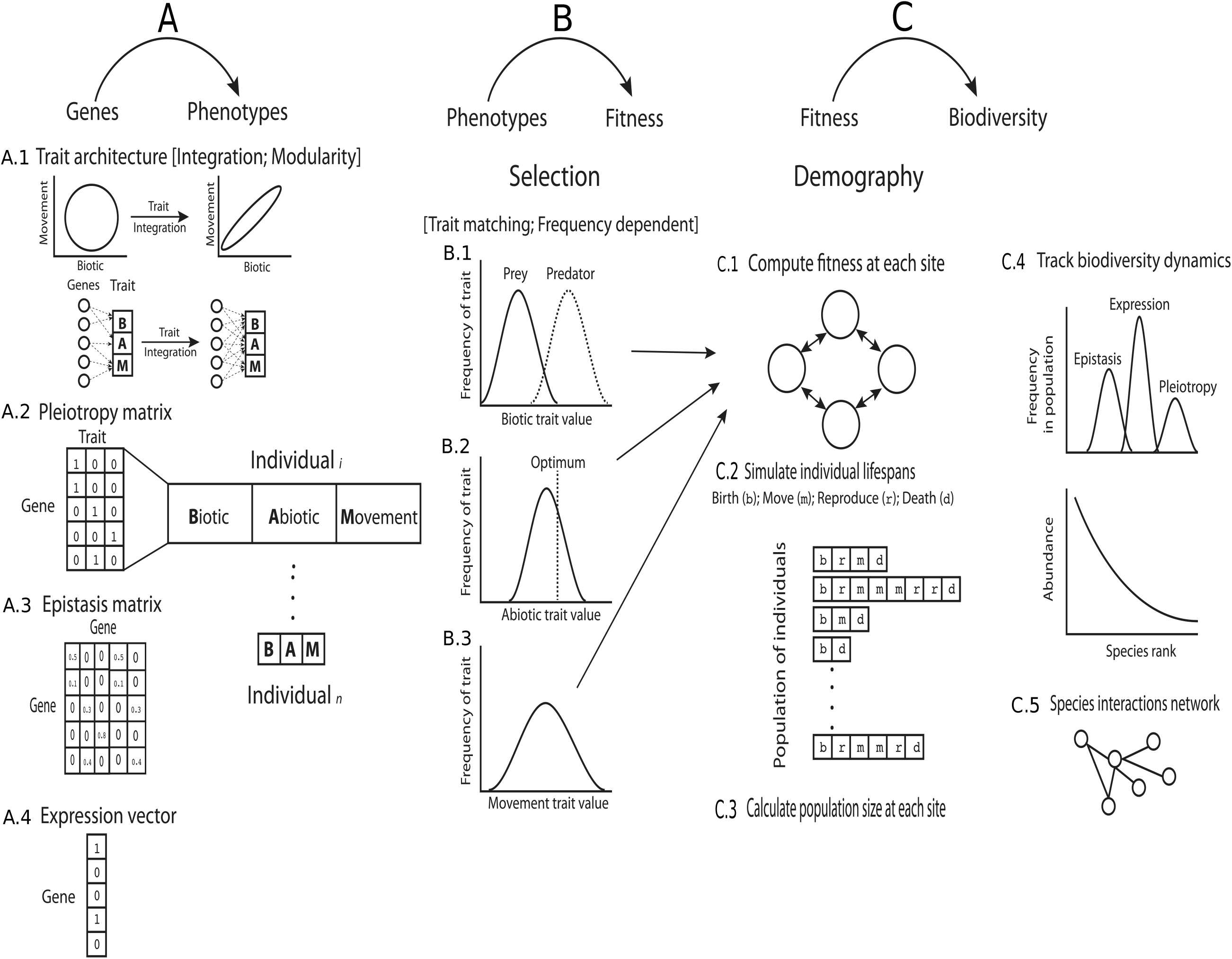
Integrating genes, phenotypes, and fitness into diversity patterns. **A) Genes to Phenotypes, B) Phenotypes to Fitness** and **C) Fitness to Biodiversity. A) Genes to Phenotypes** The two GPAs explored (**A.1**) modular, independent traits (left), and correlated, integrated traits (right). Pleiotropy (**A.2**), Epistasis (**A.3**) and the expression vector (**A.4**) determine the trait vector of each individual (vector *Z*, equation 3). **B) Phenotypes to Fitness** introduces selection accounting for (**B.1**) Biotic, (**B.2**) Abiotic and (**B.3**) Migration traits to obtain the fitness of each trait architecture. **C) Fitness to Biodiversity** computes (**C.1**) fitness of individuals at each site, (**C.2**) the life cycle of the events, death, birth, and migration and (**C.3**) the population dynamics at each site to obtain diversity metrics ranging from the distribution of trait architecture (**C.4**) to species abundances and species interaction networks (**C.5**). The figure shows the specific case for the BAM scenario: **A) Genes to Phenotypes** explores the within-species and the overall-community distribution of trait architecture following the **B) Phenotypes to Fitness** for the BAM traits. As for the general framework, the **C) Fitness to Biodiversity** computes Biodiversity metrics for the different trait architecture scenarios.

Many studies have explored GPAs with traits jointly impacted by selection or the selection on trait combinations rather than on traits in isolation to explore how they affect local and global adaptive evolution (Schluter, 1996; Wagner and Zhang, 2011; Armbruster, 2014; Svens-son et al., 2021). On the other side, many empirical and theoretical studies in developmental and evolutionary biology, as well as in molecular systems biology support the evolution of modularity in complex organisms (Callebaut and Rasskin-Gutman, 2009). Development occurs through a series of discrete modules, a modular GPA is usually composed of a high number of dimensions allowing certain parts of the organism to change without interfering with the functions of other parts (Wagner et al., 2007). Overall, the GPA to biodiversity connection might facilitate the merging of complex traits into spatially explicit models of metacommunity and food web dynamics (Figures 1 and 2) combining engines with complementary mechanisms (Box 2), thereby providing a path to expand our predictions on how the complex trait architecture will respond to fluctuations and perturbations in a rapidly changing world.

## 4 Methods

We introduce a model incorporating epistasis, pleiotropy, and gene expression to characterize the GPA of individuals and populations. Figures 1 and 2 summarize the formulation of the two GPAs explored: 1) trait modularity and 2) correlated selection (trait integration). Trait architecture takes into account polygenic traits, pleiotropy, epistasis and expression (Figure 1, layers 1 and 2, Figure 2A, and section “Genotype to phenotype architecture: Correlated selection and epistasis”). We then describe the “phenotype to fitness” section with biotic and abiotic traits influencing individual’s birth and death rates and population demography (layer 3 in Figure 1, Figure 2B, and section “Phenotypes to fitness”). The GPA contains selection acting on each trait in isolation, i.e., modularity, or on combinations of traits. We refer to selection acting on each trait independently as multivariate selection without genetic correlations, that is, a modular or independent trait architecture, and on combinations of traits as correlated architecture. The last stage in this section is the connection between fitness and species interactions in species-rich communities to evaluate the role of the two GPAs to diversity patterns (layers 4 and 5 in Figure 1, Figure 2C, and section “Fitness to Biodiversity”.)

Genetics models with epistasis have been deeply studied (Phillips, 2008; Lehner, 2011; Mackay, 2014). There have been mostly two modeling frameworks accounting for epistasis (Bank, 2022), one assuming probabilistic fitness landscapes directly mapping genotypes to fitness with a given amount of epistasis, i.e., the Rough-Mount-Fuji (RMF) model (Aita et al., 2000; Melo and Marroig, 2014), and the FGM (Fisher’s Geometric model, (Fisher, 1930), as defined by (Martin et al., 2007), and onwards) that ignores that epistasis could occur in the GP map, i.e., epistasis arises as a consequence of stabilizing selection for a phenotypic optimum. In our modeling framework, epistasis is defined as an interaction between genes, where the effect of one gene is modified by one or more other genes. We incorporate genes functioning as an intricate biological network, i.e., the product of one gene can influence the activity or expression of another. This interconnectedness means that the effect of a specific genetic variant can be dependent on the “genetic background” the other genes present. We follow with mutations affecting epistasis and then stabilizing selection for the abiotic phenotypic optima or the selection for the biotic trait shape trait distribution depending on trait architecture, i.e., modular or correlated selection where biotic and abiotic traits within each architecture can have differential contribution to fitness.

### 4.1 Genotype to phenotype architecture: pleiotropy and epistasis

Each individual’s phenotype is represented as a time-dependent vector, a function influenced by pleiotropy, epistasis, and the expression vector (refer to Figure 1 and Table 1 for a complete list of parameters and variables). The trait space of a population is a subset of N-dimensional real vectors, representing N real-valued traits. The trait architecture of each individual, within this N-dimensional space, is entirely characterized by pleiotropy, epistasis, and gene expression.

**Table 1:**
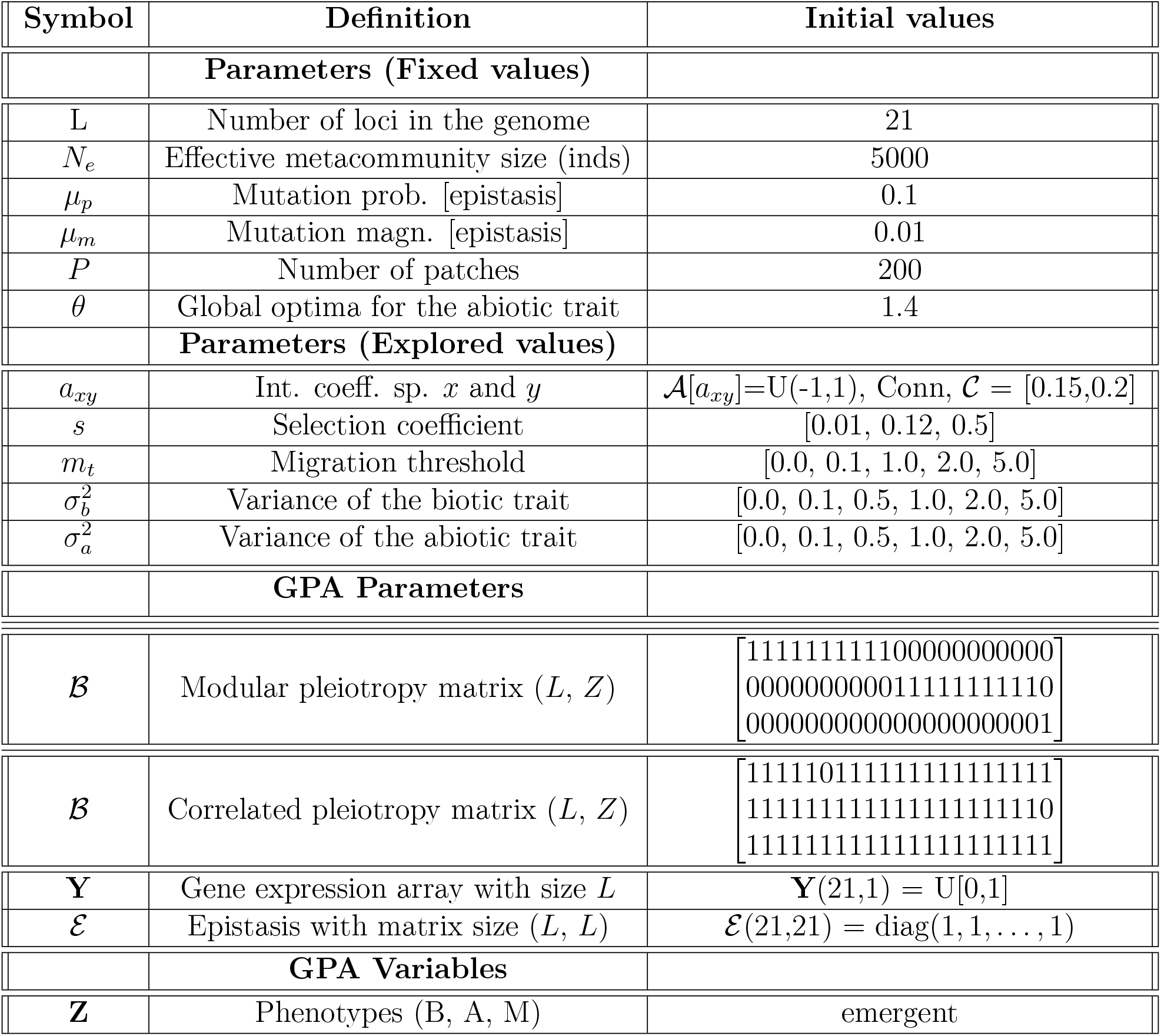
Definition of symbols and initial parameter values.

We define L as the number of haploid loci, and **L** as the vector of allelic values (or effects) at these L loci, such that **L** ∈ ℝ^*L*^. The time-dependent trait vector of each individual, **Z**(t), is thus controlled by the effects of these *m* haploid loci (represented by the vector **L**), the pleiotropy matrix, ℬ (t), the epistasis matrix, ℰ (t), and the expression vector, **Y**(t).

The effect of the loci **L** on a specific trait *i*, denoted *z*_*i*_(t), where i = 1,…, N, is initially conceived as a function:

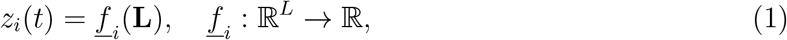

where, *f*_*i*_ represents the function for the *i*-th trait component, mapping the L loci to a single real-valued trait value.

Assuming the collective function [*f* _1_(**L**), …, *f*_*N*_ (**L**)]^*T*^ is differentiable, we can consider, as a first step, linear approximation for the genetic contribution to the traits. This linear mapping from loci effects to additive trait values is often represented by a pleiotropy matrix. We define the time-dependent pleiotropy matrix ℬ (t) as a matrix of size N × L, traits by loci, respectively, where its elements *b*_*ij*_ describe the influence of locus *j* on trait *i* (Lande and Arnold, 1983; Kauffman and Levin, 1987; Wagner, 1989; Pavlicev and Hansen, 2011; Melo and Marroig, 2014). The pleiotropy matrix, ℬ (t), with elements *b*_*ij*_, is considered a temporally-varying binary matrix where *b*_*ij*_ = 1 indicates that locus *l*_*j*_ influences trait *z*_*i*_, and *b*_*ij*_=0 indicates no influence (Melo and Marroig, 2014).

This linear approximation implies that the genetic component of the trait vector, **Z**(t), is given by:

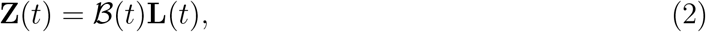

where **L**(t) represents the time-dependent allelic values of the loci. To clarify all the components, we write equation **Z**(*t*) = ℬ(*t*)**L**(*t*) in explicit matrix notation: **Z**(*t*), is the time-dependent trait vector representing the values of each trait. ℬ(*t*) is the time-dependent pleiotropy matrix connecting alleles to traits. If there are N traits and L loci, this is an N × L matrix, and **L**(*t*), represented as the time-dependent allelic values of the loci. This is a vector representing the values of the L loci. Therefore, it is an *L* × 1 matrix. Given these dimensions, the equation **Z**(*t*) = ℬ(*t*)**L**(*t*) in matrix multiplication is given by *N* × 1 = (*N* × *L*) (*L* × 1).

For clarity, let us represent the elements of each matrix and vector at a given time *t* as

- **Z**(*t*) has elements *z*_*i*_(*t*) for *i* = 1, 2, …, *N*.
- ℬ (*t*) has elements *B*_*ij*_(*t*) for *i* = 1, 2, …, *N* and *j* = 1, 2, …, *L*.
- **L**(*t*) has elements *l*_*j*_(*t*) for *j* = 1, 2, …, *L*.

The equation **Z**(*t*) = ℬ (*t*)**L**(*t*) can be written out as:

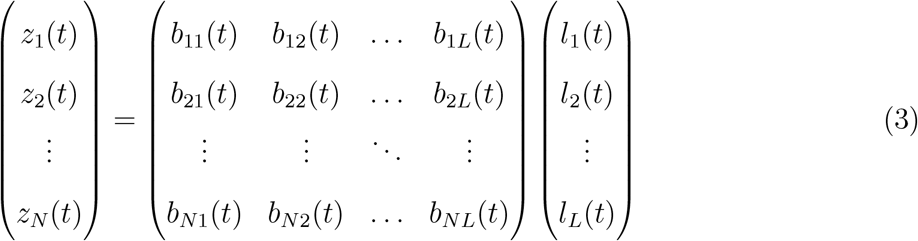

The explicit form of the equation for each trait *Z*_*i*_(*t*) is a summation. The *i*-th element of the resulting vector is the dot product of the *i*-th row of ℬ(*t*) and the column vector ℒ (*t*). Therefore, for any trait *i* (where *i* = 1, 2, …, *n*), the explicit equation is:

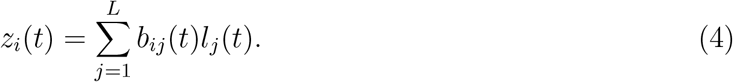

This equation states that the value of a specific trait *i* at time *t* is the sum of the products of each allelic value *l*_*j*_(*t*) and its corresponding pleiotropic effect on trait *i, b*_*ij*_(*t*). This formulation is a common way to model the genetic basis of quantitative traits.

We now add an expression component of the genes, i.e., time-dependent expression as a function of environmental states or development of individuals. The magnitude of this influence is adjusted by the expression level of each gene, which is controlled by a time-dependent expression vector, **Y**(t). We introduce the vector, **G**(*t*), as the gene expression level of each locus connecting the expression vector, **Y**(*t*) and the allelic values vector **L**(*t*). The most common way to represent a direct, element-wise relationship between the time-dependent expression vector, **Y**(*t*), and the time-dependent allelic vector, **L**(*t*) is through a Hadamard (element-wise) product (○) as:

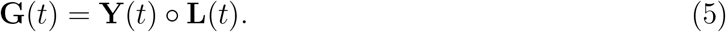

For clarity, in explicit matrix form this results as follows. With L loci, the vectors are, **G**(*t*), a vector of gene expression levels, *L* × 1, **Y**(*t*), a vector of expression states, *L* × 1, and **L**(*t*), and a vector of allelic values, *L* × 1.

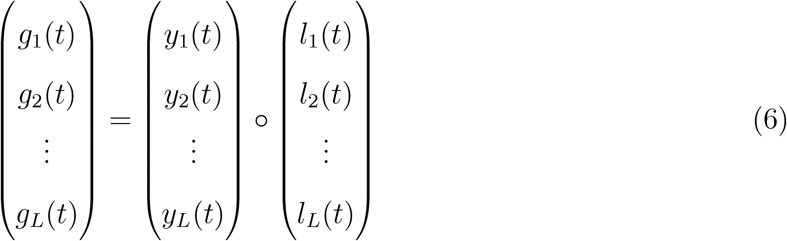

For an additional level of control to model complex traits (Baryshnikova et al., 2010; Costanzo et al., 2010), we introduce the time-dependent epistasis matrix, ℰ(t). The most direct way to introduce an *L* × *L* epistasis matrix, ℰ (*t*), is to have it operate on the gene expression vector, **G**(*t*). This operation would capture how the expression of one gene modifies the effect of another. The result of this multiplication, a new vector, would then be used later in the pleiotropy equation to determine the trait vector. We remark that the Hadamard product, **G**(*t*) = **Y**(*t*) ○ **L**(*t*), means that each element of the resulting vector **G**_*i*_(*t*) is the product of the corresponding elements from **Y**_*i*_(*t*) and **L**_*i*_(*t*), i.e., **G**_*i*_(*t*) = **Y**_*i*_(*t*) ○ **L**_*i*_(*t*). This is an explicit multiplication operation directly modeling the effect of one gene, **L**_*i*_, modified by a regulatory input, **Y**_*i*_.

For now, the equation for the time-dependent trait vector, **Z**(*t*), becomes:

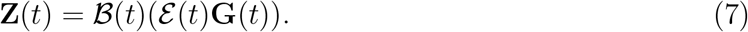

This equation can be simplified by treating the combination of the epistasis matrix and gene expression vector as a single term. The matrices and vectors involved in the whole formulation are: **Z**(*t*), *N* × 1 trait vector, ℬ(*t*), the *N* × *L* pleiotropy matrix, ℰ(*t*), the *L* × *L* epistasis matrix, and **G**(*t*): The *L* × 1 gene expression vector.

To simplify, we find the product of the epistasis matrix and the gene expression vector as

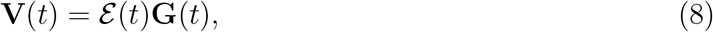

and in matrix form can be represented as:

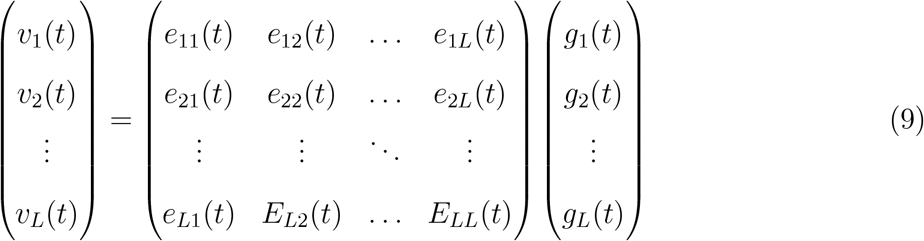

with the element *v*_*j*_(*t*) given by the summation:

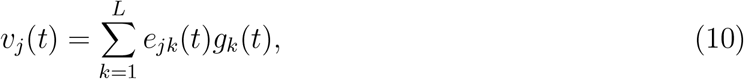

and substituting this intermediate vector back into the original trait equation we have:

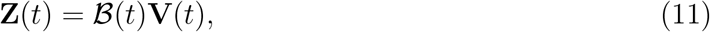

with the resulting explicit matrix form expression

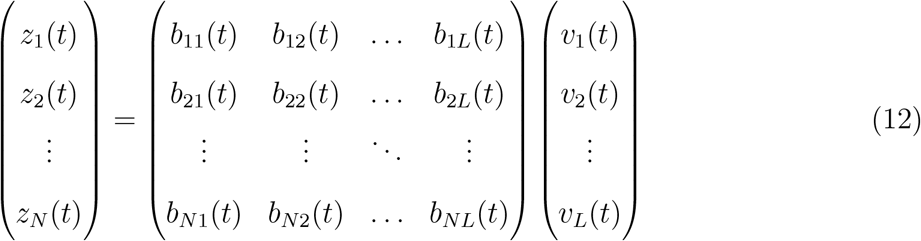

and the final explicit form for each trait *z*_*i*_(*t*) is:

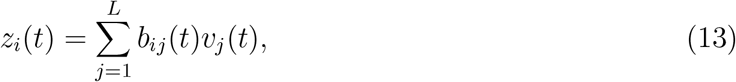

and by substituting the expression for *v*_*j*_(*t*), we get the full explicit form as:

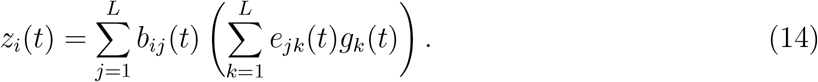

This expanded form shows that the value of each trait *i* is a function of the pleiotropic effects *b*_*ij*_(*t*), the epistatic interactions *e*_*jk*_(*t*), and the underlying gene expression levels *g*_*k*_(*t*). This represents a more complex and biologically realistic model where the effect of one gene on a trait is not only direct but can be modified by its interaction with other genes.

In order to explore how GPAs, i.e., modular or correlated affect biodiversity, we simplify the trait architecture into the following three trait classes (Figure 2): biological interactions (i.e., biotic trait: B), interactions with abiotic environmental conditions (i.e., abiotic trait: A), and movement behavior (i.e., movement and dispersal traits: M). These three types (Figure 2A) not only help us simplify the complex trait architecture of organisms (i.e., from a generalised trait vector **Z**(t) to a specific trait BAM vector). It also allows us to explore potential feedback scenarios and its connection to biodiversity patterns (Figure 2B). For example, as we describe below, we can model how the outcome of coevolutionary biotic interactions can vary spatially depending on trait architecture (e.g., modular vs correlated selection, Figure 2B) and site specific fitness optima. Furthermore, we can explicitly link how trait architecture and hierarchy between traits changes biodiversity patterns (Figure 2C).

### 4.2 Phenotypes to Fitness

Our formulation of the **Z**(t) vector allows us to derive the fitness of each individual based on their biotic, abiotic, and migration traits, i.e., the time-dependent BAM vector, as a special case of the **Z**(t) vector (Figure 2A). As we describe below we use the BAM formulation because we are interested in contrasting the two GPAs in spatially explicit landscapes. In our simulations the M trait has no specific effect on individual fitness (Figure 2). First, we compute abiotic fitness based on the mean distance of an individual’s trait value to the abiotic optimal phenotypic value for that species at that specific location *j*. The time-dependent distance of each abiotic trait to the optimum follows:

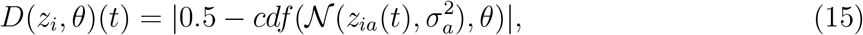

where *D*(*z*_*i*_, *θ*)(*t*) is the distance of the time-dependent abiotic trait of individual *i* to its abiotic global optimum, *θ. θ* is set in the beginning of the simulation for each species equal for all the *j* locations, 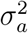 is the variance of a normal distribution that is global and species-specific, and *z*_*ia*_(*t*) is the value of the time-dependent abiotic trait *i* and *cdf* is cumulative distribution function. *D*(*z*_*i*_, *θ*) is a probability calculated by the number of standard deviations that the individual’s phenotype is different from the optimal phenotype. The variance, 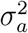, determines the severity of the deviation from the optimum, with lower values resulting in larger penalties.

The time-dependent fitness of the abiotic trait of individual *i* is then given by:

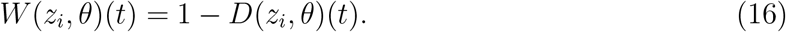

We model an individual’s biotic phenotype based on the interactions with other individuals from the same and/or difference species. Interactions between individuals depend on species-level interaction strength and sign, as well as on their individual biotic phenotypes. Two individuals’ similarity is determined by calculating the average distance for every pair of phenotypes using the following approximation:

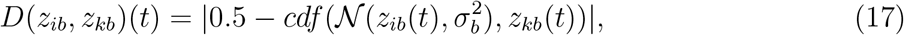

where *D*(*z*_*ib*_, *z*_*kb*_)(*t*) is the distance between individual time-dependent biotic phenotypes *i* and *k* and 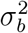 is the variance of the distribution. The larger the 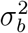, the less significant are small phenotypic differences between the two individuals. If two individuals interact, their fitness increases or decreases depending on the nature of their species-species interactions, meaning that the fitness effect of an interaction depends on the biology of each species. The time-dependent strength of an interaction is a function of species-species coefficient and the phenotypic distance between the two individuals

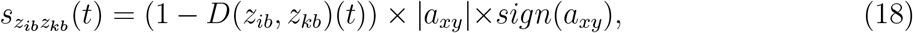

where 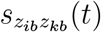 is the interaction strength between individuals *i* and *j, D*(*z*_*ib*_, *z*_*kb*_)(*t*) is the time-dependent phenotypic distance between the two individuals and *a*_*xy*_ is the interaction strength between species *x* and *y*, where individuals *i* and *k* belong to species *x* and *y*, respectively.

This “species level coefficient” refers to the community matrix 𝒜 determining trophic position of each species. *sign* returns 1 or -1 as the sign of the interaction, i.e., increases or decreases the fitness of the two individuals, respectively. We remark we do not calculate biotic fitness for each individual because that would require calculating the biotic distance for all pairs each in a site. Instead, we first let the simulation determine which two individuals actually encounter each other, and then calculate the biotic distance between these two individuals. Individual *i*’s fitness is then updated after an interaction with individual *k* as

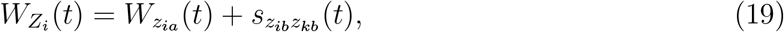

and the fitness function takes into account the selection coefficient as

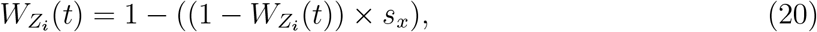

where *s*_*x*_ is the selection coefficient of individual *i* of species *x* acting on all the traits of the individual. Individual’s genotype, i.e., gene expression, pleiotropy and epistasis, map to its phenotype which is visible to selection. Overall, our formulation allows to explore different GPAs (Figure 2A, correlated selection and modularity). Different selection regimes are also allowed, e.g. varying strength of coevolutionary selection and spatially variation in fitness optima, from local variation to global optima. These processes together with migration dynamics all collectively affect trait evolution of resource and consumer species (Figure 2B-C).

### 4.3 Fitness to Biodiversity

We consider a landscape with a network of locations connected by migration and inhabited by resource and consumer populations comprising several individuals each. Biotic, abiotic and migration trait values, the BAM scheme, are sampled from eqs. 1 to 20. Population dynamics of resource and consumer species is driven by the trait- and temporal-dependent fitness functions using the BAM scheme. The procedure to connect the GPAs to demography follows individuals reproducing, dying and migrating in this order at each time step. Below we introduce the sections of our mutation-epistasis-GPA-migration model with overlapping generations: 1) demographic cycles; 2) migration, local and global optima; 3) reproduction; and 4) GPA and landscape scenarios. To account for the role of GPAs on diversity patterns, random mutations of different magnitude occur only in the epistasis matrix. Total number of individuals in the ecosystem, *N*_*e*_, is kept constant at 5,000 individuals, and this selection scheme results in effective population size equal to census population size. We run the simulations with no environmental noise, *ϵ*.

#### 4.3.1 Demographic cycles

Our modeling framework follows a sequence of events, where each sequence is repeated per time step, completing a non-zero sum dynamics cycle, i.e., number of births, deaths and migration numbers differ under the dynamics of overlapping generations. Since traits can have fitness consequences or not, we have two categories of traits: Biotic and abiotic traits with fitness consequences and migration without fitness consequences. In the GPA with BAM spatially explicit modeling we choose a biotic, an abiotic and a migration trait for each individual. Each cycle is composed of the following events in this order: aging, older individuals die with higher probability, global abiotic optima determining the abiotic component of fitness, and biotic, driven by interactions with other individuals/species in each location. All these processes are modeled on top of genomics processes described along eqs. 1 to 20, i.e., pleiotropy, mutation driven epistasis and expression all represented as the trait vector **Z**(t) (See Table 1 for definition of parameters and GPA variables).

#### 4.3.2 Migration, local and global optima

We model migration by considering a random landscape with a network of connected patches each inhabited by resource and consumer species with different interaction types. Before considering migration, the simulation checks if the migration trait is higher than a migration threshold, *m*_*t*_. If the individual satisfies the threshold criteria, it evaluates nearby sites within its visual range randomly selecting a destination. Individual fitness is updated based on the abiotic and biotic conditions at the new site. Note that in the simulations we do not impose a top-down migration rate to the species, there is no migration at that specific time step if none individual satisfies the migration threshold. If most individuals have a migration trait value above the threshold, migration rate increases, otherwise, decreases. Consequently, migration is indirectly selected to occur. We have used a migration threshold value for all the species.

#### 4.3.3 Ecological network with multiple interaction types

Unlike abiotic optima, which are set at the beginning of the simulation and remain constant across all sites, the biotic optima are not predetermined and vary for each individual at each local site. What matters for biotic traits is their relative distance to other individuals, both within and across species. This context-dependent approach acknowledges that local optima for the biotic trait differ for each species, depending on the time-varying composition of organisms in a given site. This is in line with the concept of local adaptation, where the optimal trait values are influenced by the distribution of biotic factors, such as prey and predators, competition, etc, in a specific site (Endler, 1995; Armbruster and Wege, 2018). The strength of this context-dependency is modulated by the species-species community matrix, *A*[*a*_*xy*_], a property of each species with any interaction type, i.e., the coefficients, *a*_*xy*_, can be positive and negative numbers (See Table 1).

#### 4.3.4 Reproduction

We consider a simple mating scheme to avoid the impact of assortative or disassortative mating and recombination on the genotype-to-phenotype architecture and biodiversity patterns. Haploid individuals at reproductive age produce identical offspring to themselves, except that the epistasis matrix of the offspring will mutate based on the mutation and magnitude probability (Table 1). The number of offspring is a random number from a Poisson distribution with a mean equal to the species’ growth rate times the fitness of the individual.

#### 4.3.5 GPA landscape scenarios

We explore two types of GPA: modular and correlational selection. The modular scenario takes into account independent traits, i.e., biotic and abiotic traits contributing independently to fitness (Figure 2A, modular GPA, Table 1). Correlational selection incorporates interactions between the biotic and the abiotic traits (Figure 2A, trait integration, Table 1). To understand how biodiversity patterns emerge under these two scenarios, local adaptation to abiotic and biotic factors can be viewed as a balance between several opposing forces. The degree of local adaptation depends on (i) the correlation between biotic and abiotic traits, (ii) the strength of spatially divergent natural selection acting on each trait, that is, the intensity of local abiotic and biotic selective pressures that drive genomic and phenotypic divergence, and (iii) migration, which counteracts this divergence. In contrast, global adaptation occurs when spatially distributed populations evolve a trait that is advantageous across all environments encountered by the species (our abiotic optimum follows this principle). Global adaptation of the abiotic trait is therefore defined at the phenotypic level, where all populations experience selection toward a shared global optimum. In both cases, the dynamics of global adaptation approximate those expected for a single population under directional selection, but are modified by spatial structure (Yeaman, 2022) and, in our model, by the additional effects of biotic interactions and interspecific interaction coefficients.

#### 4.3.6 Simulations

We run 1044 simulations along a selection, biotic and abiotic variance and migration gradient each for 500 generations, using parameter values outlined in Table 1 for both modular and correlational selection GPAs. We compute a decision tree and the migration rate, abiotic and biotic coefficients, and the selection coefficient distributions to explore the effect of parameter values combinations to diversity difference between the modular and the correlated GPAs (Figures 3-8).

**Figure 3.**
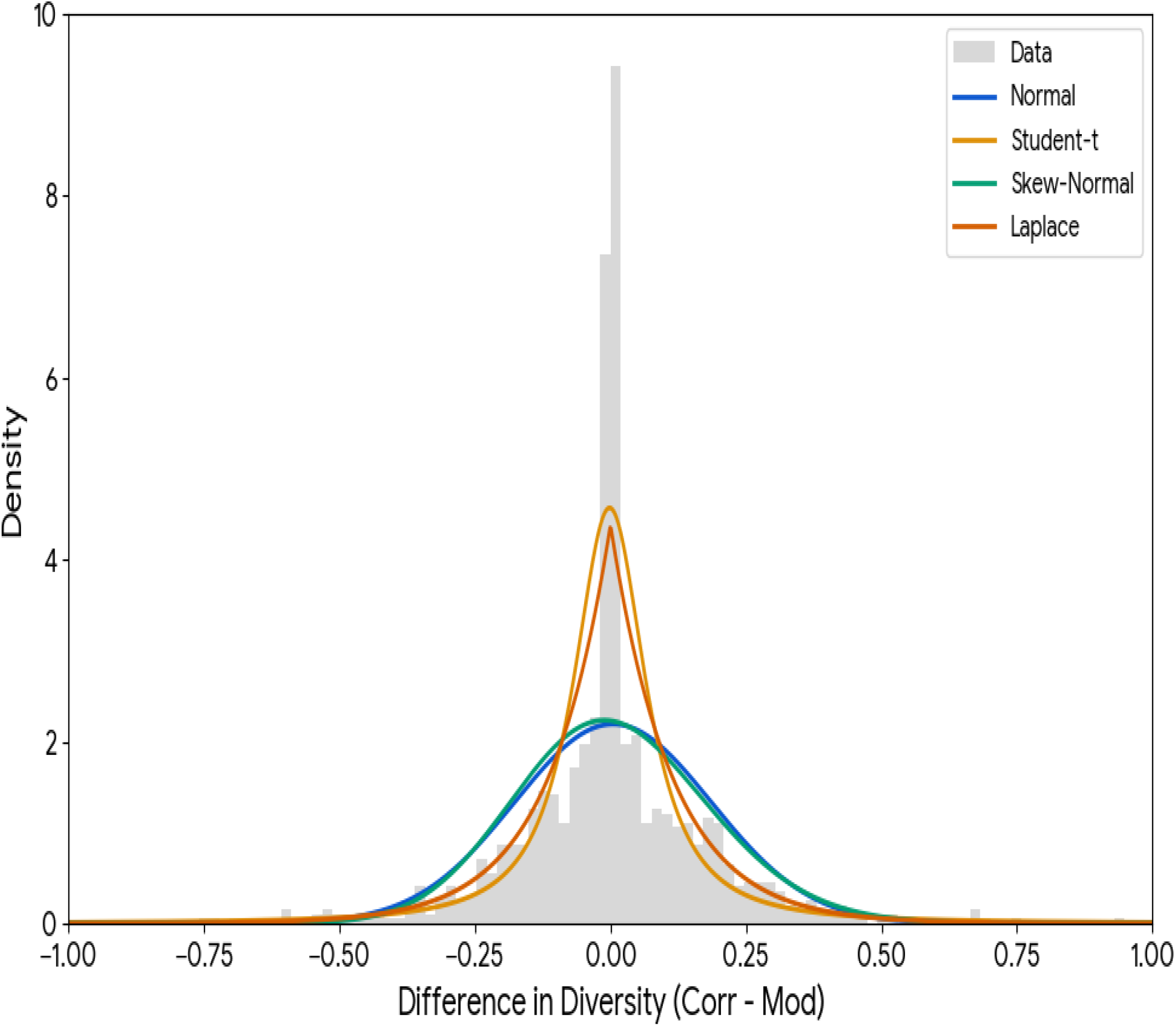
Distribution of GPA diversity difference. Histogram for the diversity difference between the correlated and the modular GPAs (grey bars) for all the simulation runs. Based on the statistical properties of all simulated data the high kurtosis (heavy tails) and significant departure from normality, the most appropriate distribution for a robust Bayesian analysis is the Laplace (Red, Laplace: Log-Likelihood = 498.42, AIC = -992.85) and the Student’s t-distribution (Orange, Log-Likelihood = 462.47, AIC = -918.94) with Normal (Blue, Log-Likelihood = 297.35, AIC = -590.69) and Skew-Normal (Green, Log-Likelihood = 308.69, AIC = -611.39) with higher AIC values.

## 5 Results

The analysis of the diversity-difference between the two GPAs with the mutation-selection-epistasis-migration model (Figures 1-2) reveals that the difference in diversity between GPAs is not a product of a single parameter, but rather an emergent outcome of a hierarchy of influences. Selection and migration consistently emerge as the “master levers” driving the diversity patterns in the correlated and the modular GPAs, respectively. While acts as the primary filter, determining the absolute magnitude of the diversity gap between the correlated and the modular GPA, serves as the directional switch that determines which architecture ultimately yields higher diversity. These results are based on the diversity-difference distribution, calculated as the difference in diversity between the correlated and the modular GPA, accounting for the same parameter combination. Simulation outputs were analyzed based on the GPA difference in diversity using Bayesian comparisons (Figure 3), the impact of selection strength on GPA diversity (Figure 4) and the Architectural Effect Intensity (Figure 5), and a GPA outlier analysis to map the impact of the selection-migration interaction landscape on diversity difference between the two GPAs (Figures 6-8).

**Figure 4.**
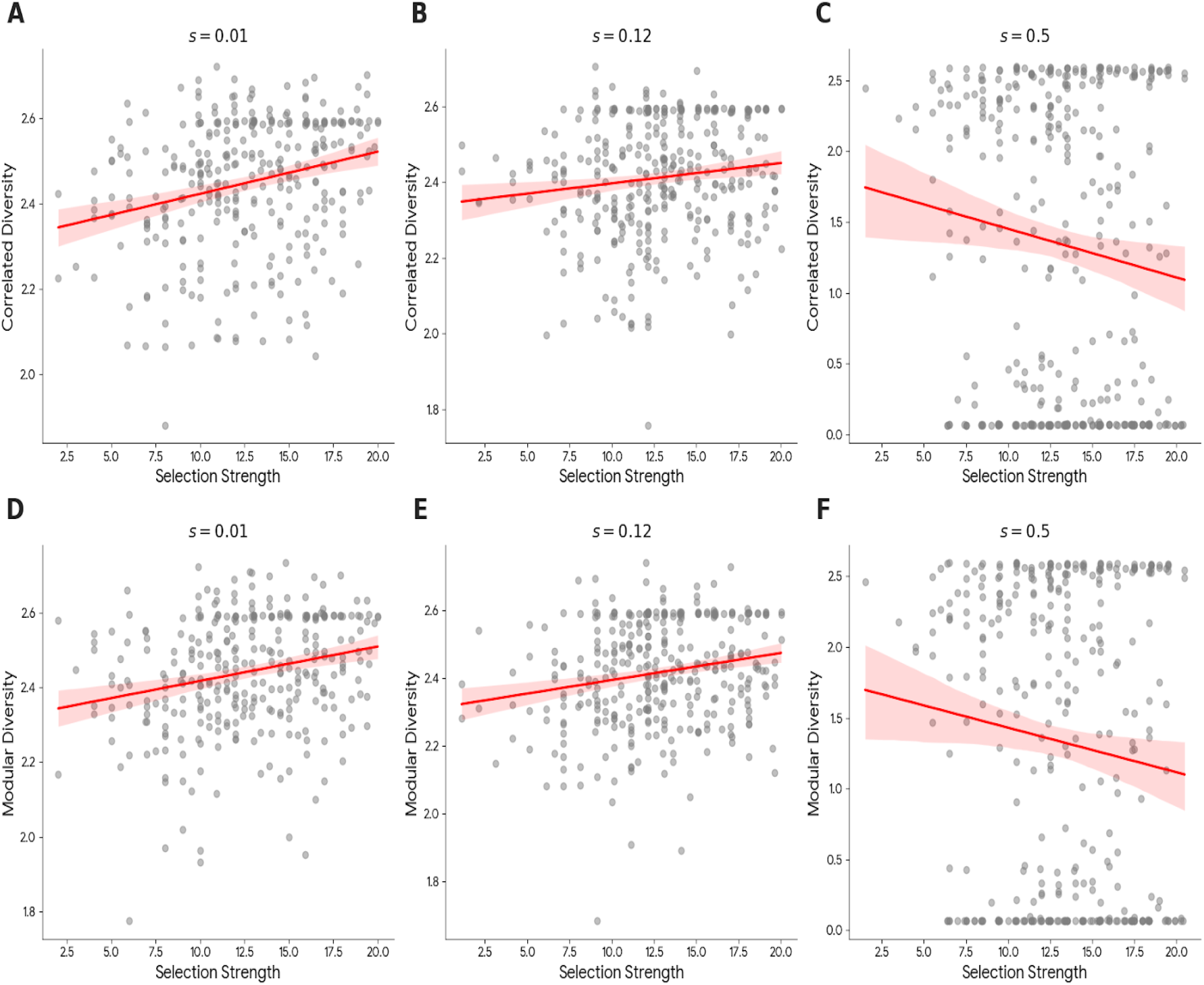
Impact of selection strength on GPA diversity. Comparison of correlated (A–C) and modular (D–F) diversity across selection gradients. At low selection coefficients (*s* equal to 0.01 and 0.12), diversity correlates positively with *s*. However, at high selection (*s* equal to 0.5), the relationship flips to a statistically significant negative correlation, indicating that intense selection begins to constrain diversity by favoring a narrow range of optimal phenotypes.

**Figure 5.**
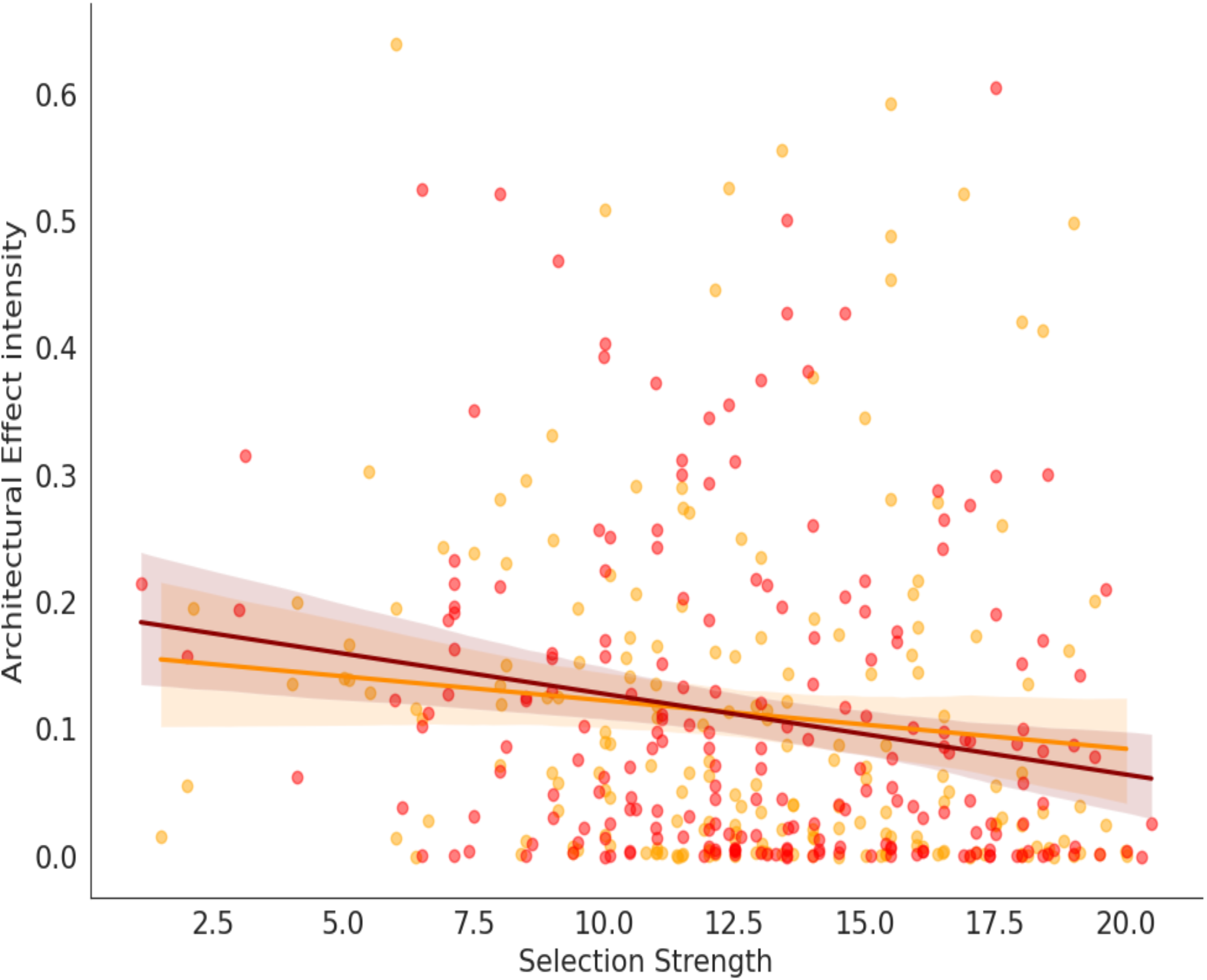
Selection strength and architectural effect intensity under varying gene flow. The absolute difference in diversity (Architectural Effect Intensity) as a function of selection and migration. Under high migration (*m* equal to 5), increasing selection strength significantly reduces architectural intensity (red, r=-0.2, p < 0.01), as gene flow homogenizes the landscape. In the absence of migration (orange), the disparity between architectures becomes less predictable (orange, r=0.11,p > 0.1), suggesting that local drift and biotic and abiotic filtering dominate.

**Figure 6.**
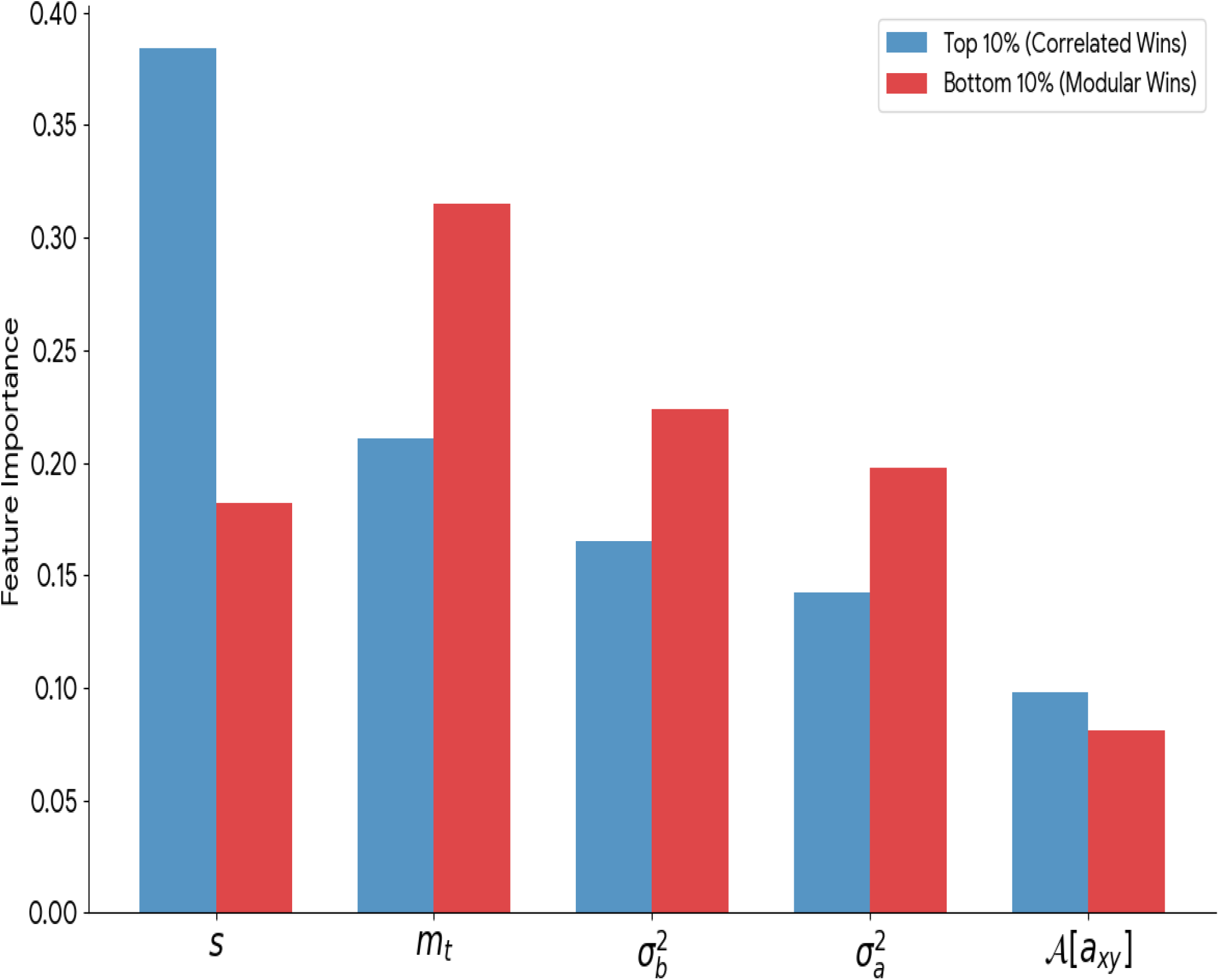
Feature importance for GPA advantage. Random Forest analysis of factors driving extreme diversity differences (Top and Bottom 10%). Correlated Advantage (Blue): Selection is the primary engine (0.384), followed by migration and biotic and abiotic variances. Modular Advantage (Red): Migration is the dominant driver (0.315), followed by biotic and abiotic variances. This highlights selection as a driver of magnitude and migration as a driver of direction.

The Bayesian comparison between correlated and modular GPA shows that neither their means nor their variances significantly differ under either the Normal or the Gamma model (Figure 3). Based on the statistical properties of the 1044 simulations, the high kurtosis (heavy tails) and significant departure from normality, the most appropriate distribution from the Bayesian analysis are the Laplace (Red, Log-Likelihood = 498.42, AIC = -992.85) and the Student’s t-distribution (Orange, Log-Likelihood = 462.47, AIC = -918.94) (Figure 3).

To explore how selection strength impacts GPA diversity, we have conducted independent analysis for each value of selection, *s* equal to 0.01, 0.12, and 0.5 for the correlated and the modular GPA using the “Distance-to-Maximum” method for selection strength, calculated as

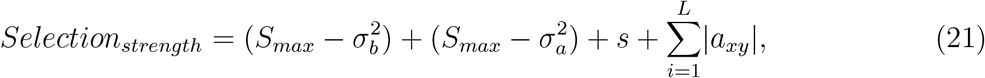

where subtracting from the maximum observed selection strength value, *S*_*max*_, accounts by the fact that larger biotic and abiotic variance represent lower selection (see Method). Similarly, the sum of the absolute value of all the links in the community matrix, 𝒜[*a*_*xy*_] accounts for the total interaction impact from the species interaction. As selection strength increases from low to medium values, both Correlated and Modular Diversity tend to increase (Correlated Div., s= 0.01, r = 0.252 and s=0.1, 0.13, p < 0.001, and Modular Div., s= 0.01, r = 0.236 and s=0.1, 0.196, p < 0.001) (Figure 4 A-B for Correlated Diversity, and C-D for Modular Diversity). For selection equal to 0.5 the relationship flips regardless the GPA. There is a statistically significant negative correlation, where higher selection strength is associated with lower diversity for both the correlated and the modular GPA (Correlated Div. s= 0.5, r = -0.124, p < 0.05, and Modular Div. s= 0.5, r = -0.114 p < 0.05.) (Figure 4 C for Correlated Diversity and F for Modular Diversity.)

The impact of selection strength on the Architectural Effect Intensity Migration plays a role as a primary gatekeeper. To decipher the impact of selection strength on the Architectural Effect Intensity we calculated the absolute difference between the correlated and the modular diversity (Figure 5). For a fixed selection coefficient, s equal to 0.5, migration becomes the most significant factor in determining how much GPAs matters (Figure 5, red, migration equal to 5, r = -0.2, p < 0.001, and orange equal to zero, r = -0.1, p = 0.12). This suggests that the spatial connectivity of populations in food webs determines the “visibility” of modular vs. correlated benefits with increasing selection strength. Therefore when migration is high, there is a statistically significant negative relationship where increasing selection strength reduces the intensity of the architectural effect. For low gene flow, the magnitude of the disparity between GPAs is less predictably tied to the aggregated selection strength index. In summary, gene flow acts as a “buffer”, making the diversity outcomes more responsive to the overriding influence of extreme selection pressures.

To gain a deeper understanding of the “extreme” cases across the entire simulation outputs, we performed a random forest analysis on the top and bottom 10% of the data, the tails of the distribution in (Figure 3) to identify what specifically drives the most significant differences between the GPAs. In such tails, there might be a shift in diversity by as much as one unit, that is nearly 50% of the mean diversity. By focusing on the extremes, we can distinguish between conditions that favor Correlated GPA (top 10%) and those that favor Modular GPA (bottom 10%). The order of importance for the correlated GPA predicting much larger diversity than the modular GPA (top 10%) starting by the selection coefficient (0.384) migration threshold (0.211), biotic variance (0.165), abiotic variance (0.142), and the interaction matrix (0.098). This suggests that high selection pressure is the primary engine that allows the correlated architecture to pull ahead of the modular one significantly. On the other side, the order for the modular GPA predicting much higher diversity than the correlated GPA (Bottom 10%) starts with the migration threshold (0.315), the biotic variance (0.224), the abiotic variance (0.198), the selection (0.182) and the interaction matrix (0.081) (Figure 6.) When modular GPA predicts higher diversity the situation is more complex. Migration takes over as the primary driver, indicating that the relative success of modularity is more dependent on population connectivity and gene flow than on the raw magnitude of selection alone.

Figure 6 also shows the drivers of diversity difference between the two GPAs are asymmetric. High selection coefficients are the primary driver when correlated GPA significantly outperform modular ones. Conversely, when modular architectures take a significant lead, migration is the most influential factor. In addition, biotic and abiotic variance are more critical in the modular GPA tail than in the correlated tail. This suggests that modular GPA may rely more heavily on specific environmental or biotic interaction regimes to gain a large advantage in diversity. The interaction matrix has the lowest relative importance in both tails, but it remains a consistent background factor, contributing roughly 8-10% to the model’s ability to classify these extreme cases. In summary, the random forest on intensity (magnitude) confirmed that selection strength dictates if there will be a large difference, but the migration threshold and biotic and abiotic conditions determine which GPA actually pulls ahead. When we isolate the extremes of the diversity-difference distribution (the top and bottom 10%), a striking asymmetry in architectural success becomes apparent. Correlated GPA achieve their greatest diversity advantage under regimes of high selection pressure, where trait correlations provide a deterministic edge in navigating fitness peaks. Conversely, the Modular advantage is significantly more context-dependent. Its success is less about the magnitude of selection and more about the spatial connectivity of the food web and the specific balance of the biotic and abiotic trait coefficients.

We have generated a Partial Dependence Plot (PDP) to examine the simultaneous effect of selection (s) and migration threshold (*m*_*t*_) on the probability that the correlated (Figure 7) or modular (Figure 8) GPA yields a significantly higher diversity, defined as being in the top (bottom) 10% of diversity difference values. The PDP can be considered as a GPA outlier analysis to map a selection-migration interaction landscape affecting GPA diversity to reinforce the distinct properties of GPAs on diversity. The correlated GPA landscape shows selection dominance. As selection increases, there is a clear shift in the probability of correlated advantage. High selection pressure acts as a filter that favors correlated GPA, regardless of the baseline diversity. The migration side is non-linear. In low migration regimes, the GPA effect is more sensitive to small changes in selection. As migration increases, the “probability pockets” become more stable, suggesting that gene flow acts as a buffer against architectural differences between correlated and modular GPAs. In summary, the correlated GPA outperform the modular one at the intersection of high selection and moderate migration.

**Figure 7.**
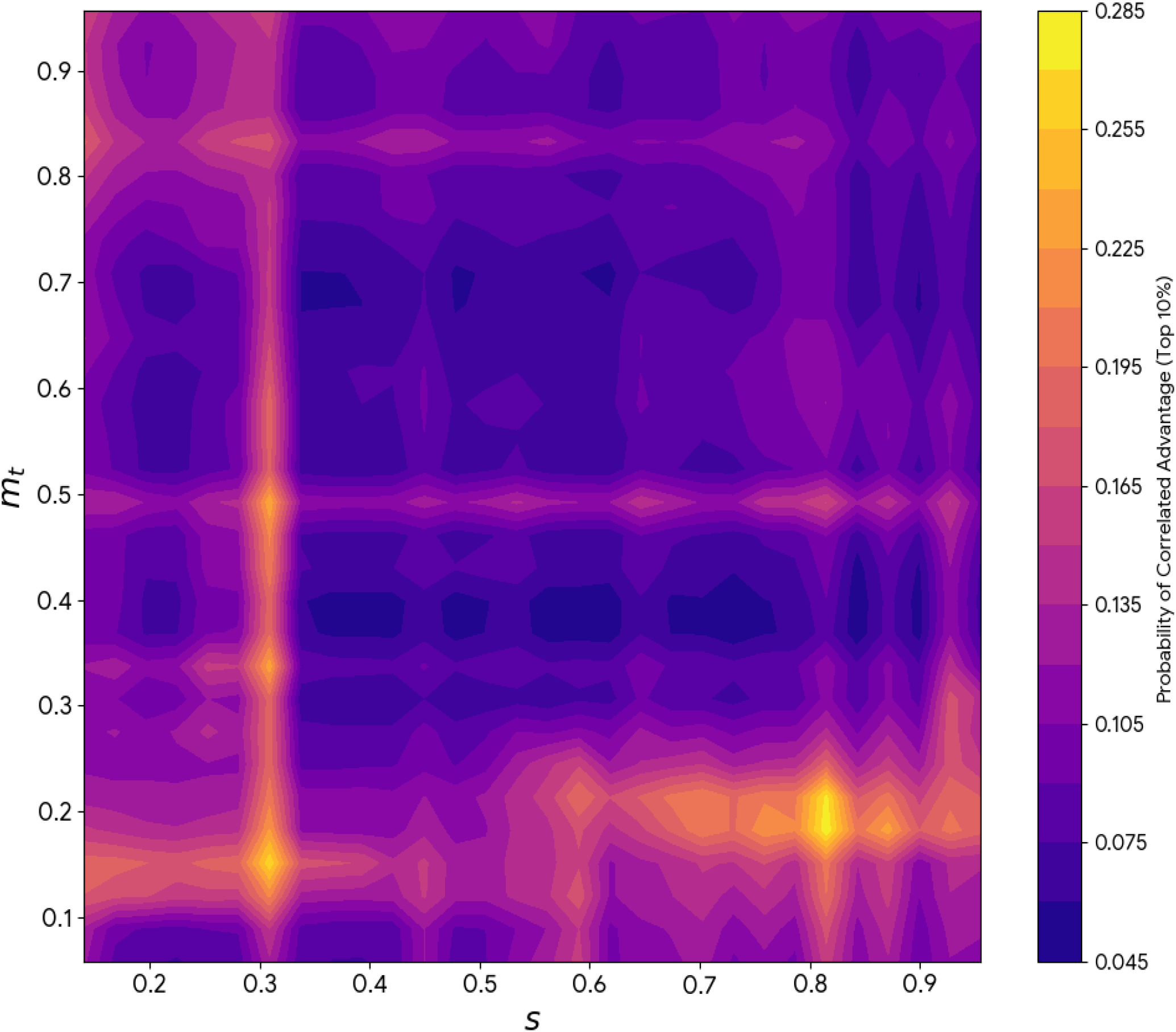
Probability landscape of correlated GPA advantage. Partial dependence plot showing the simultaneous effect of selection and migration on the probability of correlated GPA winning in diversity. The landscape is characterized by selection dominance; as selection increases, the correlated GPA provides a consistent diversity advantage regardless of baseline connectivity.

**Figure 8.**
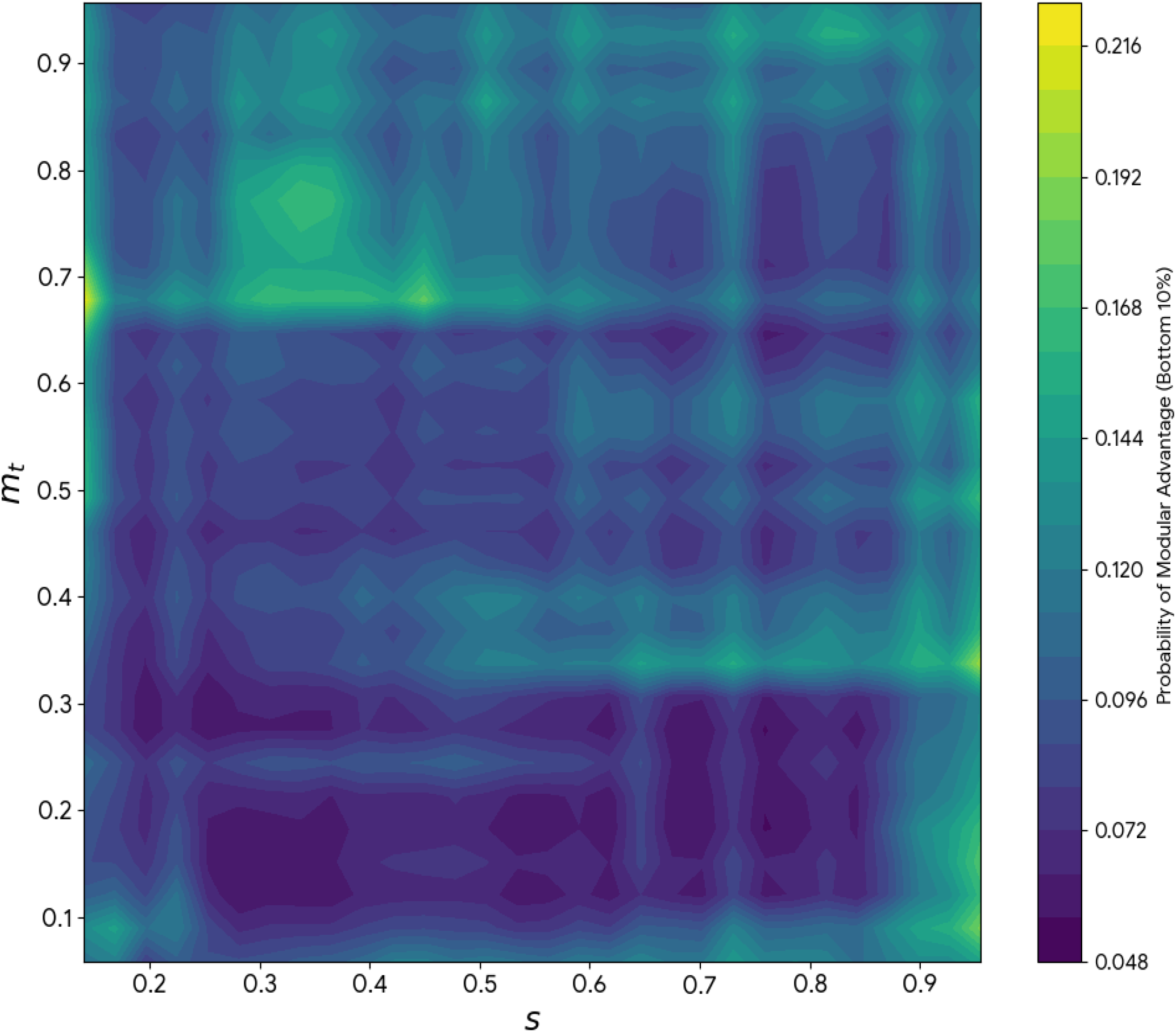
Probability landscape of modular GPA advantage. Partial dependence plot showing the probability of modular GPA advantage. Unlike the correlated landscape, modularity shows a complex, non-linear dependence on migration. Modular advantage peaks in distinct regions of the parameter space, specifically where high migration rates allow the decoupling of trait modules to navigate spatially conflicting selective pressures more effectively.

The modular GPA landscape shows a more complex dependence on migration. In several regions, increasing migration shifts the likelihood of a modular GPA outperforming a correlated one, highlighting its role as a key driver for higher diversity in the modular GPA. The probability of modular advantage in diversity often peaks in different parts of the landscape compared to the correlated GPA. This confirms our previous finding that while high selection generally separates the two GPAs, the specific differences in diversity between the two GPA is determined by the interplay with community and food web connectivity. In summary, modular GPA may rely on specific balances between local adaptation and gene flow to maximize their diversity relative to correlated GPA.

## 6 Discussion

Decades of research have mapped global patterns of genetic and species diversity (Leigh et al., 2021; Gaston, 2000; Antonovics, 1992; Vellend, 2005), yet the functional “wiring” that connects these levels remains a significant frontier. While recent advances acknowledge that selective optima and genetic covariances dictate the pace of evolution (Walsh and Blows, 2009; do O and Whitlock, 2023), most multispecies models still overlook the underlying genotype-phenotype architecture (GPA) Melián et al. (2011); Moya-Laraño et al. (2014); Hagen et al. (2021); Overcast et al. (2021). Many studies have also focused on correlated selection or the selection on trait combinations rather than on multiple traits in isolation (Schluter, 1996; Wagner and Zhang, 2011; Svensson et al., 2021; Armbruster and Wege, 2018). Many experiments in systems biology, on the other side, have shown the evolution of modularity, where traits can evolve in isolation of one another (Wagner et al., 2007; Callebaut and Rasskin-Gutman, 2009). By employing a “minimalist model” of complex trait architecture, we demonstrate that biodiversity is not merely a product of selection magnitude, but of how that selection is filtered through complex modular versus correlated trait architectures. Our study also provide a likelihood approximation (Box 1) and a synthesis of the existing digital engines (Box 2) to combine the parametrization of complex trait models from GWAS data to explore biodiversity in a more integrated digital ecosystem.

Our most salient finding is the emergence of a hierarchy among eco-evolutionary drivers. While selection coefficient determines the intensity of the architectural effect, the migration rate and the nature of the selection (biotic vs. abiotic) dictate the direction of the advantage (Figures 5-8). Specifically, we find that correlated architectures, characterized by high pleiotropy across trait classes—predict higher diversity under regimes of low migration and moderate, balanced selection. In these “isolated” landscapes, genetic and ecological drift appear to favor the integrated trait responses of correlated GPAs. Conversely, modular architectures emerge as a better strategy for maintaining diversity under high migration and strong biotic interactions. By allowing traits to evolve in relative isolation, modularity provides the flexibility required to adapt to the conflicting selective pressures inherent in highly connected, species-rich communities. We notice that we are contrasting two GPAs, correlated selection and modularity, with a fixed covariance matrix, i.e., GPAs do not evolve during our simulations. Second, complex GPAs explicitly coupled to pleiotropy, epistasis and expression dynamics exploring much larger genomes and phenomes with nonlinear effects is still far when connecting complex trait architecture to biodiversity dynamics (Blows et al., 2015). For example, separating cause and effect in studies of the genetic covariance and the direction of evolution would require explicitly accounting for a changing covariance matrix, genetic association studies, i.e., GWAS data, fluctuating environments and more flexible local and global optima (do O and Whitlock, 2023; Yin et al., 2024), contrary to what we have simulated in our biotic and abiotic fitness-related traits.

Many studies have shown the evolution of clusters of locally adaptive loci, i.e., outlier clusters (Yeaman, 2013), which can arise through both non-neutral and neutral processes (Nosil et al., 2017; Southcott and R., 2017). These genomics clusters may represent different underlying trait architectures. For instance, they might reflect the presence of a modular trait architecture, where strongly interacting and partially isolated genes each influence a distinct subset of traits. Alternatively, such clusters could result from strong pleiotropy, where a small number of genomic regions exert broad influence, affecting many traits simultaneously, both within and across modules. The two types of trait architecture explored in this study, modular and correlated selection, could be connected to the genomic clustering patterns. As these clusters evolve, the genomic clusters and “outlier” regions suggests that architectures themselves are under selection (Nosil et al., 2017). Future iterations of this modeling framework should account for evolving covariance matrices and developmental plasticity to better capture how species “re-wire” their GPMs into different GPAs, from modular to correlated and viceversa, in response to rapid biotic and environmental change. Furthermore, the tension between local and global optima, here explored through a globally adapting biotic trait, remains a critical pivot point. Understanding how the gain or loss of modularity or correlated selection occurs across a species’ range will be essential for predicting evolutionary rescue versus extinction in multifarious perturbation scenarios.

Connecting GPAs to eco-evo-devo in species-rich communities requires refinement of the dynamics introduced in this study. First, many empirical studies have shown epistasis and pleiotropy can rapidly change across generations. The covariance matrix encoding trait architecture likely changes across generations, impacting population dynamics. Second, developmental processes change as well as regulatory processes during the ontogeny of individuals. Accounting for the dynamics of covariance matrix and development could allow partitioning the role of biotic and abiotic traits into the ecological, developmental and evolutionary components explaining biodiversity patterns (Post and Palkovacs, 2009; Schoener, 2011). We did not account for developmental symbiosis and developmental plasticity, despite having explicit expression-regulation as a fixed vector in the simulations. Developmental symbiosis and developmental plasticity can be introduced by adding changing rates to the expression vector within individuals. These processes affect fitness through non-genomic mechanisms of selectable and heritable variation, including adaptive variation and niche construction (Gilbert et al., 2015).

An additional level of improvement when connecting GPA to biodiversity would be to consider the dynamics under local vs. global optima. Global adaptation in spatially structured populations implies a biotic and/or an abiotic trait, or the GPA as a whole, within a species that is beneficial in all environments the species encounters. Global adaptation can be defined at the phenotypic level, where all populations of a species experience selection toward the same optimum, at the trait or loci level, where a given locus or a set of loci has the highest average fitness across the range of the species and natural selection tends to favor its fixation throughout. In either case, global adaptation tends to behave approximately according to dynamics expected for a single population under directional selection, but with some modifications due to the effect of migration and spatial structure (Yeaman, 2022). Our abiotic trait is under global adaptation; the global optima is the mean abiotic trait distribution at the outset. Coupling strong selection of a globally adapting abiotic trait to correlational selection GPA might nullify the effect of many architectures from the overall dynamics, and thereby decrease intraspecific and species diversity. On the other side, unique abiotic local optima might favour the selection of different trait architectures. In summary, one of the main challenges in connecting GPAs to biodiversity dynamics is to account for more realistic local and global optima to understand how the gain and loss of GPAs within species occur when there is a tension between migration, local and global optima selection regimes.

Empirical and theoretical studies support the notion that eco-evolutionary feedbacks are expected when evolution is rapid (Fussmann et al., 2007b). Yet, the role of the structure of feedbacks when connecting ecological, developmental and evolutionary processes into rapid GPA changes are mostly unknown (King and Wilson, 1975; Carroll, 2005; Marasco and Kornblihtt, 2023). Rapid evolutionary changes might be especially common in antagonistic interactions, such as predator-prey and host-parasite systems, but also in competitive and mutualistic interactions because phenotypes within species might be constantly responding to a co-evolving biotic selective pressure (Yoshida et al., 2003; Morran et al., 2011). Feedback, however, might occur with different traits, i.e., demographic traits throughout population demography and trait evolution (Kokko and andrés, 2007), and abiotic traits between organisms and abiotic conditions (e.g. evolution by niche construction, (Matthews et al., 2014)). Ultimately, biotic and abiotic feedbacks might also be mediated by the incorporation of regulatory-expression mechanisms via developmental dynamics. Many morphological and cellular traits are the result of complex physico-chemical processes because gene actions are intimately linked to developmental interactions; thus, the regulatory-expression dynamics taking into account organismal complexity in development, might provide new information on how development, GPA and biodiversity connect each other under rapidly changing ecosystems.

Our approach enables the exploration of development effects on eco-evolutionary and coevolutionary feedbacks in multispecies dynamics, and facilitates a transition in focus from single (Lion, 2018; Govaert et al., 2019; Weber et al., 2017; Terhorst et al., 2018) to multiple traits (Alberch, 1991; Thompson et al., 2013; Salazar-Ciudad et al., 2021; Niraula et al., 2024) while allowing for biotic and abiotic selection pressures from individuals to food webs. By using a “minimalist model of complex traits” into multilayer networks, we aim to move beyond single trait models to a community-level framework accounting for the “internal” complexity of species. Extensions of this approach allows to partition the roles of development, inheritance, and environment in shaping biodiversity. Ultimately, linking complex trait architecture to the dynamics of entire assemblages provides a more robust, “mechanistically wired” path toward forecasting species’ fates in a rapidly changing world.

## 7 Acknowledgements

This work has been partly funded by the SNSF Scientific Exchanges grants IZSEZ0frm[o]–83489 and IZSEZ0frm[o]–83490 to Dr. Cecilia S. Andreazzi, Dr. Julia Astegiano and Dr. Carlos J. Melián. Thanks to Catherine Graham, Ayana Martins and Malena Sabatino for their feedback during the workshop “Coevolutionary dynamics in Metacommunities (COMETA)” organized as part of the SNSF Scientific Exchanges grants. PRG was funded by CNPq (307134/2017–2), FAPESP (2018/14809–0). PRG and CJM were funded by SNSF-Spark (CRSK-3_2_28777).

## 8 Code and data availability

All the simulations have been done in the EvoDynamics.jl julia package *EvoDynamics*.*jl* (Vahdati and Melián, 2022). Codes and data supporting results are in the public GitHub repository Cometa.

## 11 Tables and Boxes

### Box 1: GWAS data to infer GPA trait distributions

**1. The Core Model (GPA)**

Genetic-wide association data (GWAS) can be used to infer GPAs with epistasis and pleiotropy (Yin et al., 2024). To link an individual’s genotype to its *i*-th complex trait value, *z*_*i*_, we account for pleiotropy, *b*_*ij*_(*t*), the ℬ matrix, and epistasis, *e*_*jk*_(*t*), the ℰ matrix following equation 14 in the main ms. Notice that the regulation vector in equation 14, **G**(*t*), all have values equal to one:

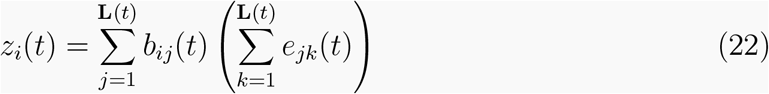

**2. GPAs Hypotheses**

We contrast the effects of two GPAs, where both the pleiotropy (ℬ) and epistasis (ℰ) matrices are structured. This means that the epistasis Matrix (ℰ) for the modular GPA has trait-specific modules (Sparse/Block-Diagonal) and interactions concentrated within modules. For the correlated GPA loci are integrated, affecting multiple traits, i.e., dense connectivity, and interactions widespread across the entire genome.

**3. Parameterization and Likelihood**

To ensure the estimation is robust to outliers and heavy tails common in complex epistatic traits, we use a Student’s *t*-distribution. The log-likelihood becomes:

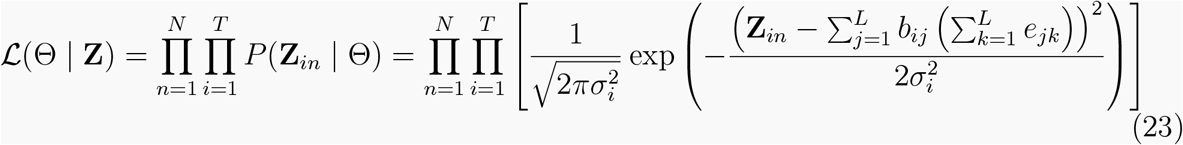

For practical optimization, finding the best parameters Θ, given the GWAS data, **Z**_*in*_, we use the Log-Likelihood ln(ℒ), as it turns the product into a sum:

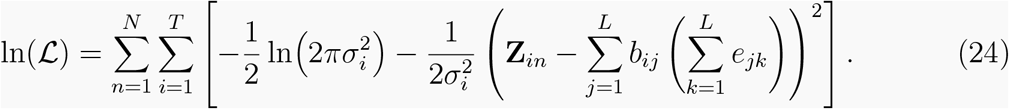

Maximizing this log-likelihood allows us to parametrize the model, i.e., find the optimal values for *b*_*ij*_ and *e*_*jk*_) using the GWAS data **Z**_*in*_ and 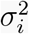 estimates. Assuming the observations across *N* individuals and *T* traits are independent, the total likelihood is the product of the individual likelihoods:

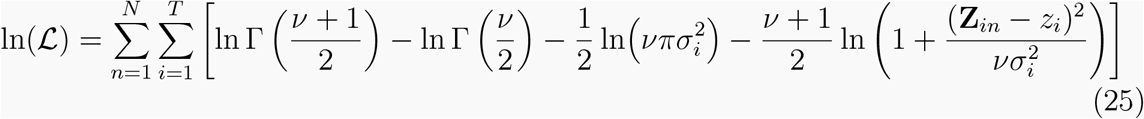

where *ν* represents the degrees of freedom and the terms 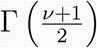 and 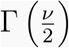 involve the Gamma function as part of the normalizing constant (or partition function). They ensure that the integral of the probability density function over the entire domain is exactly 1, which is a requirement for any valid probability distribution. This formulation prevents rare phenotypic extremes from disproportionately influencing the estimation of pleiotropic and epistatic coefficients.

**4. Simulation with population size, N, equal to 2000**

The Log-Likelihood values show the sensitivity of the metric to the correct GPA:

| Data Source (True GPA) | Candidate Model Used | Log-Likelihood ($\ln(\mathcal{L})$) |
| --- | --- | --- |
| Modular Data | Modular GPA (True) | -9668.19 |
| Modular Data | Correlated GPA (False) | -9978.52 |
| Correlated Data | Correlated GPA (True) | -9668.19 |
| Correlated Data | Modular GPA (False) | -9976.86 |

- **Architecture Discrimination:** The substantial gap (∼ 310 LL units) between the true model and the misspecified model confirms that the robust *t* Log-Likelihood is highly effective for discriminating between the Modular and Correlated GPAs.

**5. Distinct Phenotypic Distributions**

The distinct phenotypic outcomes confirm that the GPA is the primary driver of observed trait patterns.

- **Modular GPA:** Tends to produce distributions with lower variance and means closer to zero, as genetic effects are localized.
- **Correlated GPA:** Produces distributions that are more widely dispersed (higher variance) and subject to greater shifts in the mean, as the correlation effects propagate across the entire genome.

### Box 2: The emerging digital ecosystem connecting complex GPA to biodiversity dynamics

Polygenic traits have been detected for many complex traits and large genomes are being sequenced, yet characterizing the dimensionality of genomes and annotating them to the few or many loci affecting traits and their role to overall fitness is still challenging Enbody et al. (2023). Modeling polygenic traits accounting for pleiotropy, epistasis and expression-regulation gradients in the context of multiple species assemblages face scalability, mathematical tractability and robustness issues. Models with few parameters are fast, mathematically tractable and robust (e.g., stochastic recombination-mutation-migration population genetics models with epistasis are in this side), whereas those with a large number of parameters are slower, less tractable and less robust but can capture higher organismal complexity (models with full genomes accounting for pleiotropy, epistasis and regulatory processes together with varying heritability and plasticity for complex GPA scenarios are in this other side).

Trade-offs in modeling approaches make the digital ecosystem an asset to complement angles to build a more unified synthesis. There is a convergence around building individual based agents taking into account varying degrees of organismal complexity. **SLIM 4** Haller and Messer (2023) **combines explicit chromosomes and complex genomics processes with species interactions. The glads** R-package simulates GP maps with varying complexity in genomics processes and mating schemes to study islands of genomic divergence in metapopulations Quilondrán et al. (2020). **Weaver** takes explicit genomics processes connected to many traits, from lifespan to metabolic rates, to quantify the strength of species interactions in complex landscapes Moya-Laraño et al. (2014). Novel unifying engines might complement frameworks dealing with organismal complexity but have scalability issues. For example, **Gen3sis** Hagen et al. (2021), **MESS** Overcast et al. (2021), and **CDMetaPOP 2** Day et al. (2023) might be faster and can have high tractability and robustness, but the connection between high organismal complexity accounting for explicit genomes and complex traits to the macroscopic patterns of biodiversity accounting for large biogeographic regions has not been made.

A rarely explored hybrid short-cut framework might be to build on classic population genetics dealing with genomics and trait distributions, single or multiple evolving traits under the infinite loci assumption for some traits, but not for others, and from single quantitative traits to many complex traits. This approach would require contrasting basic Ornstein-Uhlenbeck processes Gardiner (1985); Hansen (1997) to more complex population genetics modeling approaches Taper and Chase (1985); M. and Kondrashov (1999) to contrast fast, scalable and robust analytical expectations with the slow, non-scalable and low robustness ones to complement the different angles Johri et al. (2022). Future developments around hybrid short-cut approaches could be merging the modular structure of the existing engines to cross-validate predictions coming from fast, scalable and robust analytical expectations with slow, non-scalable and non-robust numerical simulations. Cross-validation of the empirical patterns coming from these two angles, the simple and complex GPA connecting to contemporary and deep time and space might help to decipher the coupling between GPA, diversification and biodiversity dynamics and the evolution of ecosystem functions.

